# Are we really Bayesian? Probabilistic inference shows sub-optimal knowledge transfer

**DOI:** 10.1101/2023.04.06.535669

**Authors:** Chin-Hsuan Sophie Lin, Trang Thuy Do, Lee Unsworth, Marta I. Garrido

## Abstract

Numerous studies have found that the Bayesian framework, which formulates the optimal integration of the knowledge of the world (i.e. prior) and current sensory evidence (i.e. likelihood), captures human behaviours sufficiently well. However, there are debates regarding whether humans use precise but cognitively demanding Bayesian computations for behaviours. Across two studies, we trained participants to estimate hidden locations of a target drawn from priors with different levels of uncertainty. In each trial, scattered dots provided noisy likelihood information about the target location. Participants showed that they learned the priors and combined prior and likelihood information to infer target locations in a Bayes fashion. We then introduced a transfer condition presenting a trained prior and a likelihood that has never been put together during training. How well participants integrate this novel likelihood with their learned prior is an indicator of whether participants perform Bayesian computations. In one study, participants experienced the newly introduced likelihood, which was paired with a different prior, during training. Participants changed likelihood weighting following expected directions although the degrees of change were significantly lower than Bayes-optimal predictions. In another group, the novel likelihoods were never used during training. We found people integrated a new likelihood within (interpolation) better than the one outside (extrapolation) the range of their previous learning experience and they were quantitatively Bayes-suboptimal in both. We replicated the findings of both studies in a validation dataset. Our results showed that Bayesian behaviours may not always be achieved by a full Bayesian computation. Future studies can apply our approach to different tasks to enhance the understanding of decision-making mechanisms.

**Author summary:** Bayesian decision theory has emerged as a unified approach for capturing a wide range of behaviours under uncertainty. However, behavioural evidence supporting that humans use explicit Bayesian computation is scarce. While it has been argued that knowledge generalization should be treated as hard evidence of the use of Bayesian strategies, results from previous work were inconclusive. Here, we develop a marker that effectively quantifies how well humans transfer learned priors to a new scenario. Our marker can be applied to various tasks and thus can provide a path forwarding the understanding of psychological and biological underpinnings of inferential behaviours.

## Introduction

How should we make sensible decisions using uncertain and ambiguous information? This is a major challenge we face in daily life. Bayesian decision theory (BDT) posits that probabilistically rational decisions can be reached by integrating knowledge about the environment (i.e. prior) together with current sensory inputs (i.e. likelihood) based on their respective reliabilities (1). Evidence has shown that across various perceptual and cognitive tasks (2–8) behaviours are close to Bayes-optimal. However, whether Bayesian behaviours are computed by the brain using the Bayes rule remains hotly debated (9–11). It has been argued that given the complexity of Bayesian computations, the brain may well use simpler approximations to achieve Bayes fashioned behaviours (11,12). Reports showing behaviours that are qualitatively Bayesian (i.e., following the reliability-based weighting principle) but fall short of Bayes-optimal (review see 13) cast doubt upon the ideal Bayesian observer theory. Understanding the exact strategies behind decisions under uncertainty is important given how ubiquitous uncertainty is in every real-life decision (14).

It has been proposed that instead of using how well performance matches Bayes-optimal prediction to define an ideal “Bayesian observer”, we should use a “generalization” criterion to evaluate whether the brain indeed makes inference in a Bayesian way (15). While people can also learn a Bayes optimal policy by trial and error, these kinds of brute-force learners need to re-learn the policy whenever a change occurs in a learned task. As a result, they would likely fall short of the mark right after the change. On the contrary, a truly Bayesian agent that fully learns and flexibly accesses the components of the BDT will generalize efficiently by integrating new changes in the likelihood to any known priors (16). Generalization can be experimentally confirmed if a learned prior is transferred to novel conditions. This process has been coined as *Bayesian transfer* (15). Importantly, when a divergence from optimal transfer happens, its pattern can reveal the cognitive policy used. To further illustrate, policies that emerge from knowing the underlying statistical information of the world provide generalization to unseen data beyond the context of past observations. On the other hand, policies that do not entail a model that well represents the real world will likely fail to generalize (15) or only show “generalization” behaviours when within the context of past observations (17,18). Transfer is thus more informative than optimality matching in advancing our understanding of decision making strategies in the brain.

A few studies have investigated Bayesian transfer in perception. In tasks where strong priors from years of experience are used (e.g. inferring object locations given auditory and visual cues such as locating a vehicle from its image and engine sounds in daily life; ref 19) or when prior probability distributions are visually presented (20,21), people instantly integrate changes of prior-likelihood pairs, which has been interpreted as successful transfer. However, where two embedded location priors were acquired through trial-by-trial feedback during a task (22), participants failed to show a robust transfer effect when a new likelihood was introduced. The authors of this study (22) concluded that in their task, the complexity of two priors and the cognitive demand of acquiring them simultaneously may have prompted the use of a computationally less expensive non-Bayesian strategy, and hence a weak transfer. These studies suggested that the cognitive load of learning and maintaining priors might be a deciding factor on whether people use a Bayesian strategy.

Besides prior complexity, the characteristics of sensory inputs (likelihoods) encountered during transfer also affect transfer effectiveness. Studies observed successful transfer when a likelihood distribution has either been acquired through lifelong experience (19,23) (in both studies the manipulated senory reliabiltity is visual contrast) or used during learning (20,21) (in both studies the manipulated sensory reliability is how likely a cue location is the true target location and all the cues used in the transfer tasks were presented during learning albeit with a prior not used in the transfer). Conversely, Kiryakova and colleagues (22) evaluated transfer by introducing a likelihood that participants had never experienced when they learned the location task and found trivial transfer. This difference touches on a fundamental issue about inferential behaviours. It is often considered that the human mind excels at applying knowledge to unseen data and untrained tasks (24,25). Many argued that structured world knowledge which can represent the generative processes of physical stimuli is needed to enable such powerful generalization. The ability of the Bayesian model to explain how humans build abstract knowledge from sensory inputs is thus strongly appealing to neuroscientists (26,27). However, an alternative hypothesis is that the seemingly powerful generalization is an illusion caused by the hardware (i.e. our brains) that has long evolved to fit the world we live in well (24,28) and been optimised with extensive training data encountered during lifetime (18). Based on this hypothesis, nearly perfect generalization in response to novel likelihoods only occur when these likelihoods are physical features that are either well represented in the brain as a result of natural selection (such as luminance, ref 19) , or within the context of past experience (17). On the other hand, generalizing outside the natural selection process or living experience would be suboptimal. Overall results from the abovementioned transfer studies seem to support predictions by this “bounded optimal” hypothesis. However, as have been described, the weak transfer in (22) could have simply resulted from multiple prior learning so more studies with proper designs that can discriminate effects from each factor are needed.

“Successful transfer” has been defined by behaviours being close to Bayes-optimal in novel conditions according to objective prior/likelihood distributions. However, research has repeatedly shown that humans can behave qualitatively Bayesian and yet represent prior/likelihood uncertainty differently from objective parameters (13,21,29). Importantly, in these studies, Bayesian models still explain behaviours better than non-Bayesian models, meaning that neither can imperfect prior/likelihood weighting exclude the use of Bayesian strategy, nor can close-to-Bayes-optimal be taken as evidence for transfer. In fact, Kiryakova and colleagues (22) acknowledged that a lack of transfer in their study may be due to a small effect size caused by biased uncertainty estimation. Therefore, examining transfer using a proper mathematically operationalised criterion that can separate influence of imperfect prior/likelihood estimation from suboptimal transfer is needed before we can draw any conclusions about the effect of cognitive demand on the use of Bayesian strategies or consider any implications of transferring outside training data.

In this study, we propose a mathematical definition of transfer that is prior/likelihood independent and provides a quantification of knowledge generalization. We applied a visual-spatial task (**Fig 1.A&B**) adapted from (30) in which the location of a hidden target is sampled from prior distributions (with low or high variance that participants learn from location feedback) and scattered dots provide noisy likelihood information about target location each trial (with variance manipulated using dot dispersion). Human participants could use both prior and likelihood to estimate positions of hidden visual targets. We report two experiments. The first asked whether the use of Bayesian strategies depends on cognitive loads. To do so, we compared transfer performance between sequential and simultaneous learning of two priors (**Fig 1.C**). Based on past studies, we hypothesised that transfer would be worse in simultaneous learning due to its higher cognitive demand. The second experiment asked whether knowledge transfer depends on generalizing within (i.e., interpolation) or outside learning (i.e., extrapolation) conditions (**Fig 1.D**). Based on past studies we hypothesised that transfer would be close to optimal in the interpolation but suboptimal in the extrapolation condition. Further, in response to the replication crisis (31), for both experiments, we present data from a discovery and an independent validation dataset.

**Fig 1.**
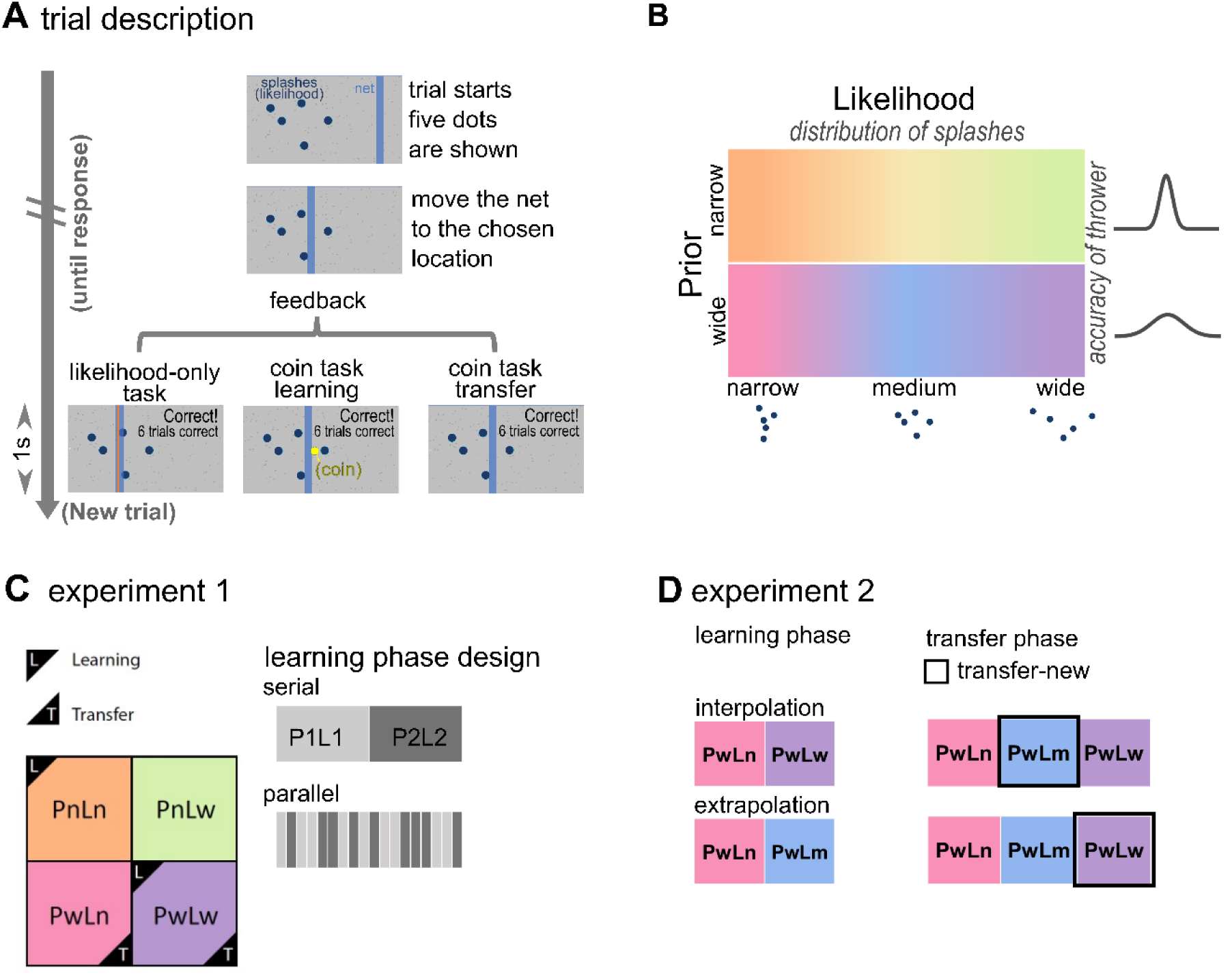
Experimental design A. Trial description in likelihood-only task, coin task - learning and coin task - transfer phases. In the beginning of each trial, participants saw 5 dots which represented splashes caused by a thrower throwing a coin into the pond (grey screen). Participants moved a net (blue vertical bar) to the position where they think the centre of the splashes (likelihood-only task), or the coin (coin task) is. After participants submitted a response, feedback information was displayed for 1 second on the screen. The next trial then started automatically. Feedback information differed between tasks and phases. In the likelihood-only task, the horizontal position of the real centre of splashes was displayed as an orange vertical line. In the learning phase of the coin task, the true coin position was displayed as a yellow dot every trial. An accumulated score was displayed when participants hit the coin. In the transfer phase, only an accumulated score was displayed. **B** Task design: Likelihood uncertainty was manipulated through the dot dispersion. Prior uncertainty was manipulated through the accuracy of the thrower, which participants learned from coin position feedback during the learning phase of coin task. Specific prior and likelihood combinations of the experiment 1 and 2 are explained in detail as follows. **C** Design of experiment 1. There were two priors (narrow Pn σ = .025 and wide Pw σ = .085) and two likelihoods (narrow Ln σ = .06 and wide Lw σ = .15). Participants underwent two orthogonal prior/likelihood combinations in the learning phase and then one prior coupling with both likelihoods in the transfer phase. In the figure example, the learning conditions are PnLn and PwLw (boxes with “L” ticks) while the transfer conditions are PwLn and PwLw (boxes with “T” ticks), with PwLn being the new combination of the transfer phase. For the serial group, combinations in the learning phase were delivered block-wise. For the parallel group, combinations in the learning phase were delivered in an interspersed way. Trials were always administered in an interspersed way in the transfer phase. **D** Design of experiment 2. In the learning phase, participants underwent combinations having one prior (wide Pw σ = .085) paired with two out of three (narrow Ln σ = .024, medium Lm σ = .06, wide Lw σ = .15) likelihoods. During learning, the interpolation group experienced PwLn and PwLw trials while the extrapolation group experienced PwLn and PwLm trials. All participants then undertook PwLn, PwLm and PwLw trials in the transfer phase. For the interpolation group, PwLm was the new combination. For the extrapolation group, PwLw was the new combination All trials were administered in an interspersed way.

## Results

### Experiment 1

Linear mixed models were used for slope (**Fig 3.A & B**) because they were not normally distributed. We used likelihood ratio tests to compare linear mixed-effect models that were designed to delineate the effects of prior, likelihood, and learning group on slopes. ANOVA was used for transfer score (**Fig 3.C & D**).

#### Learning phase

##### Discovery set

A model with the prior and likelihood fixed effects explained slope data best. We showed that participants were qualitatively Bayesian (**Fig 3.A**), i.e. weighting according to the reliability of prior (linear mixed model *p* = 6.27 X 10^-14^) and likelihood (linear mixed model *p* = 8.80 X 10^-16^) but they gave more weights to the likelihood than Bayesian optimal observers would have (one tailed sign rank test compared to optimal slope *p* <.00001 except the *PwLn* combination). No statistical evidence showed that the two cognitive load groups behaved differently in the learning phase. Models which included group as a fixed effect did not fit the data better than models which did not and slope values between the two groups were not significantly different (Wilcoxon rank sum test *p* = .84).

##### Validation set

(**Fig 3.C**) A linear mixed model that includes prior, likelihood and cognitive load group and their interactions as fixed-effect terms explained the data best (log likelihood ratio 51.48 compared to a model with prior and likelihood and their interaction as the fixed effect, *p* < .001). There was a significant three-way interaction (*p* = 9.10 X 10^-4^) as well as a group-likelihood (*p* = 7.33 X 10^-07^) and a prior-likelihood (*p* = 4.15 X10^-5^) interaction. Further analyses found that the statistical result was explained by a bigger slope difference between *PnLn* and *PnLw* trials in the serial group, as compared to those of the parallel group (*PnLn* in serial median±iqr = .85±.42, *PnLw* in serial median±iqr = .25±.30; *PnLn* in parallel median±iqr = .78 ±.36, and *PnLw* in parallel median±iqr = .66±.43). Importantly, despite the numerical difference, both groups were qualitatively Bayesian, i.e. weighting according to the reliability of prior (prior effect of linear mixed model: serial group *p* = 0.01; parallel group *p* = 1.50 X 10^-5^) and likelihood (likelihood effect of linear mixed model: serial group *p* = 3.57 X 10^-18^; parallel group *p* = 2.28 X 10^-8^). Like the discovery set, in both cognitive load groups, participants gave more weights to the likelihood than Bayesian optimal observers would have (one tailed sign rank test compared to optimal slope *p* < .00001 in all prior/likelihood combinations except the *PwLn* trials).

#### Transfer phase

##### Discovery set

We separated transfer phase data into “old” and “new” trial types. “Old” indicates those prior/likelihood combinations that individual participants had experienced during the learning phase while “new” indicates those combinations that were introduced in the transfer phase. Linear mixed models constructed to comparing the “old” combinations in the learning and transfer phase found that the model with prior and likelihood fixed effects explained slope data best. We showed that reliability-based weighting maintained in the “transfer-old” trials (linear mixed model; prior *p* = 7.45 X 10^-3^; likelihood *p* = 1.60 X 10^-4^, **Fig 3.A**), meaning that behaviours remained Bayesian after the removal of location feedback. This was further supported by a non-significant phase effect (Wilcoxon signed rank test *p* = .17). We then inspected the “transfer-new” trials. No group effect was observed (model including group effect v.s. model not including group effect: log likelihood ratio = 2.05, *p* = .22). There was a significant difference in slope between prior types but not between likelihood types (linear mixed model; prior *p* = .01, likelihood *p* = .32) implying participants may not have transferred the knowledge in a way that ideal Bayesian observer should have (**Fig 3.A**). However, it is also obvious that from **Fig 3.A**, slopes varied widely between participants even in the learning phase. This noise in the data could have greatly diminished the power of detecting transfer. To remove slope variabilities caused by subjective prior and likelihood estimations, we computed the “transfer score” (***ts***) (**Fig 2.B**, also see the **Methods** section). As intended, transfer scores indeed did not differ between prior and likelihood conditions (three-way ANOVA; prior main effect *p* = .98, evidence of main effect BF_10_ = .29; likelihood main effect *p* = .99, evidence of main effect BF_10_ = .14; **S1 Fig.A**); supporting the “transfer score” a prior/likelihood-invariant and more valid measure of transfer. However, there was an interaction between group and prior (*p* = .03). Post-hoc analysis showed that transfer score of narrow prior trials in the parallel group were lower than wide prior trials in the parallel group though not statistically significant (***ts_parallel-Pn_*** mean ± SE = 0.15 ± .12, ***ts_parallel-Pw_*** mean ± SE = 0.56 ± .10; *p* = .07). We found that there was significant (i.e. larger than zero; right tailed t-test; *p* = 7.16 X 10^-11^) but suboptimal (i.e. smaller than one; left tailed t-test *p* = 9.92 X 10^-16^) Bayesian transfer. Transfer scores in the serial group were numerically higher than those in the parallel group but not statistically different (***ts_serial_*** mean±SE = 0.50±.08, ***ts_parallel_*** mean±SE = 0. 35±.08; two sampled t-test *p* = 0.21; **Fig 3.C**).

**Fig 2.**
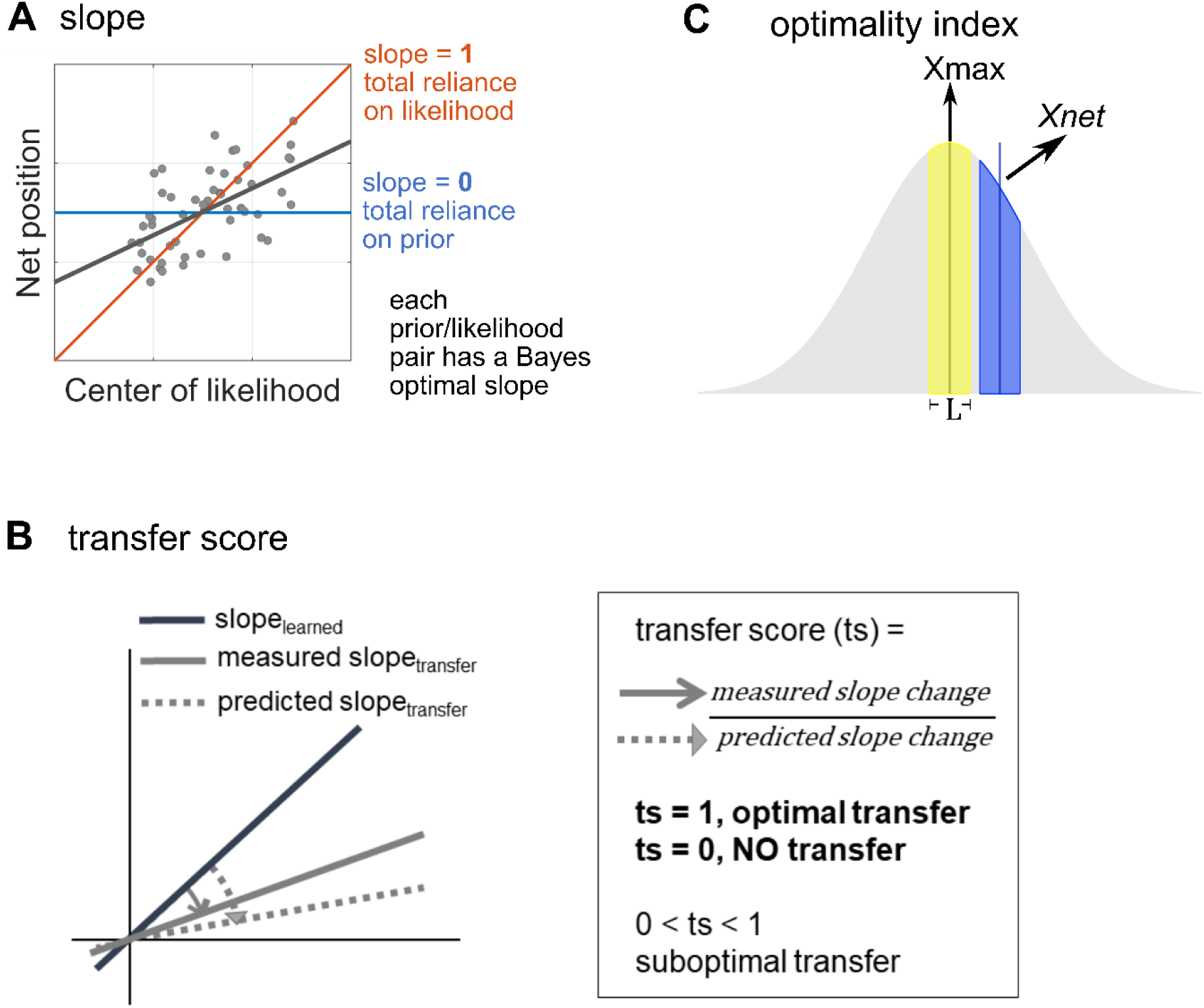
visualisation of quantitative performance measures including slope (A), transfer score (B) and optimality index (C). **A** Slope in the coin task was calculated by linearly regressing participants’ estimated coin position over the centre of the splashes. The values of slope vary between 0 and 1. A higher slope means a higher weighting on likelihood, with 1 meaning total reliance on likelihood and 0 meaning no reliance on likelihood. **B** A transfer score was calculated by normalising measured slope change by predicted slope change based on subjective prior and likelihood estimations. A transfer score of 1 means transferring following an optimal Bayesian observer model. A transfer score equals or smaller than 0 means no transfer. **C** optimality index. For each trial given the true posterior, we can compute the probability that a coin would be within the net from the chosen position (X_net_), i.e., the success probability. We defined the optimality index for a trial as the success probability normalised by the maximal success probability. In the figure, this equals to the blue area divided by the yellow area. Note that here for visualisation purpose only, there is no overlap between the two areas, which may not and does not have to be the case in real measurement.

##### Validation set

A linear mixed models with prior and likelihood fixed effects constructed to compare the “old” combinations between the learning and transfer phases explained the slope data best. We showed that reliability-based weighting maintained in the “transfer-old” trials (linear mixed model; prior *p* = 3.43 X 10^-6^; likelihood *p* = 0.002, **Fig 3.B**), meaning that slope remained qualitatively Bayesian and did not differ after the removal of location feedback (“old” combinations of the learning and transfer phases, Wilcoxon signed rank test *p* = 0.78). We then inspected the “transfer-new” trials. Again, no group effect was observed. There was a significant difference in sensory weighting between prior types but not between likelihood types (linear mixed model; prior *p* = 2.22 X 10^-05^, likelihood *p* = .94; **Fig 3.B**), as has been found in the discovery set.

**Fig 3.**
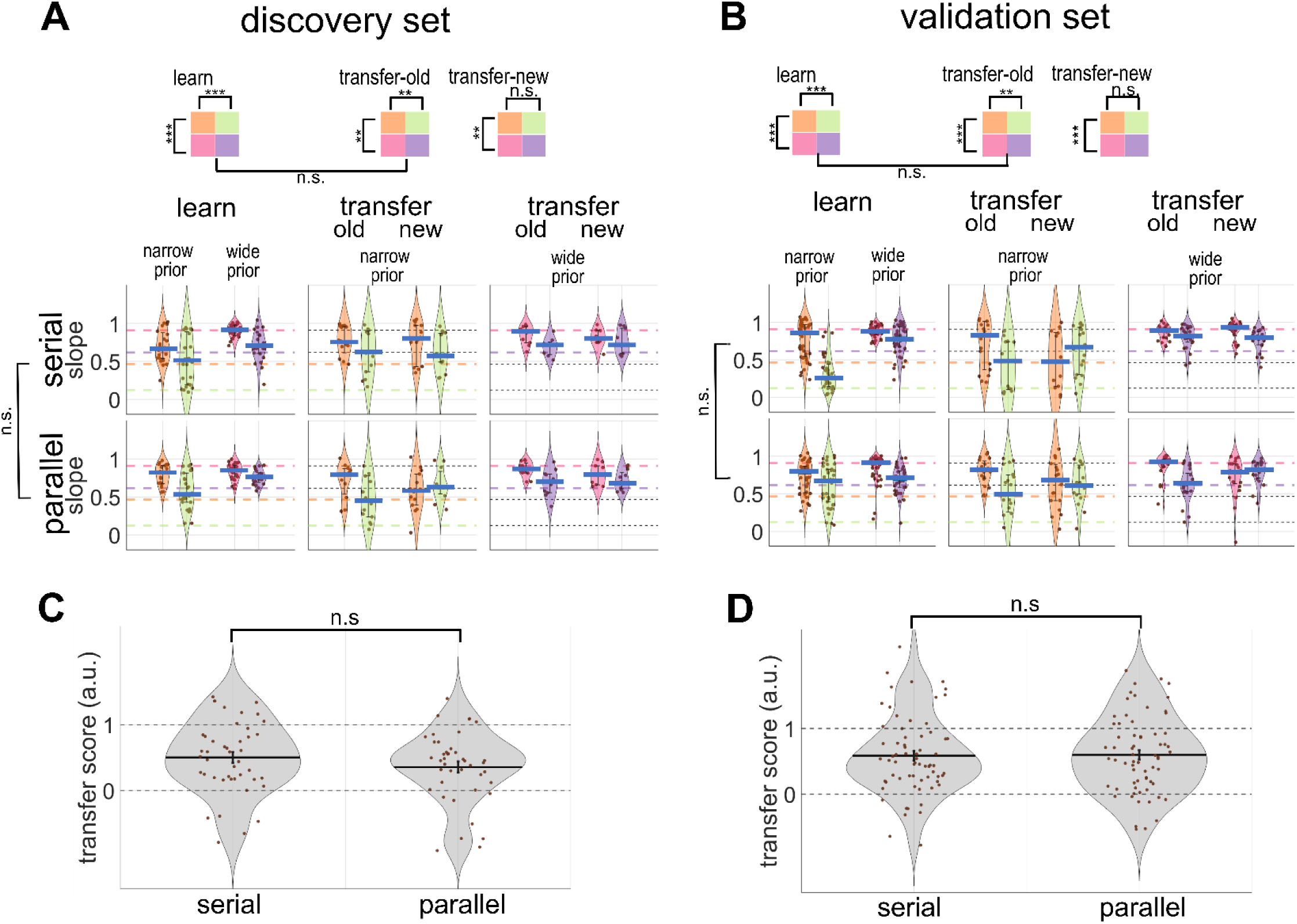
Slope and transfer score in the experiment 1. slope A Discovery & B Validation set. Violin plots show the distribution of slopes in the learning, transfer-old and transfer-new trials of experiment 1, separated by prior, likelihood and cognitive load group (upper serial, lower parallel). The central blue line shows the median. The error bar represents the interquartile range. Filled dots represent each participant. Each prior/likelihood combination is represented by orange PnLn, green PnLw, pink PwLn, and violet PwLw. Bayes-optimal values are presented as coloured dashed lines, with colours corresponding to prior/likelihood types. **transfer score B Discovery & D validation sets** The distribution of transfer scores in the serial and parallel groups. The central black line shows the mean. The error bar represents the standard error. ** p<= .01; ***p<=.001; n.s. non-significant

Transfer scores did not differ between prior and likelihood conditions (three-way ANOVA: prior main effect *p* = .11, evidence of main effect BF_10_ = .35; likelihood main effect *p* = .90, evidence of main effect BF_10_ = .14; supporting information **S1 Fig. B**). Nor was any interaction or group main effect (group *p* = .55) found. We again identified significant (i.e. larger than zero; right tailed t-test *p* = 5.68 X 10^-22^) but suboptimal (i.e. smaller than one; left tailed t-test *p* = 2.36 X 10^-13^) Bayesian transfer. Transfer score was .58±.07 (mean±SE) in the serial group and .60±.07 (mean±SE) in the parallel group (two sampled t-test *p* = 0.91, **Fig 3.D**).

### Experiment 2

Slope data in the Experiment 2 were not normally distributed either and linear mixed models were used while ANOVA was used for transfer score.

#### Learning phase

##### Discovery set

A full model which includes the likelihood and group effects, and their interaction explained the slope data best (log likelihood ratio as compared to a model with the likelihood and group effect but no interaction 12.91, *p* = .01). A significant likelihood-group interaction was found (*p* =2.50 X 10^-4^). Pos-Hoc analysis showed that slopes were qualitatively Bayesian for both interpolation (median ± iqr = .98 ± .05 for *PwLn* and median ± iqr = .86 ± .18 for *PwLw*; Wilcoxon sign rank test *p* = 4.54 X 10^-7^) and extrapolation (median ± iqr = .99 ± .04 for *PwLn* and median ± iqr = .97 ± .05 for *PwLm*; Wilcoxon sign rank test *p* = .005) groups, i.e. participants decreased sensory weighting as the variance of likelihood became bigger. We concluded that the interaction was caused by a larger disparity between prior/likelihood pairs in the interpolation group than which in the extrapolation group. Like experiment 1, participants however relied on likelihoods more than Bayesian optimal observers would have in the medium and wide likelihood conditions (one tailed sign rank test compared to optimal *slope* interpolation – *PwLw p* = .0001, extrapolation – *PwLm p* = 2.40 X 10^-7^, **Fig 4.A**).

**Fig 4.**
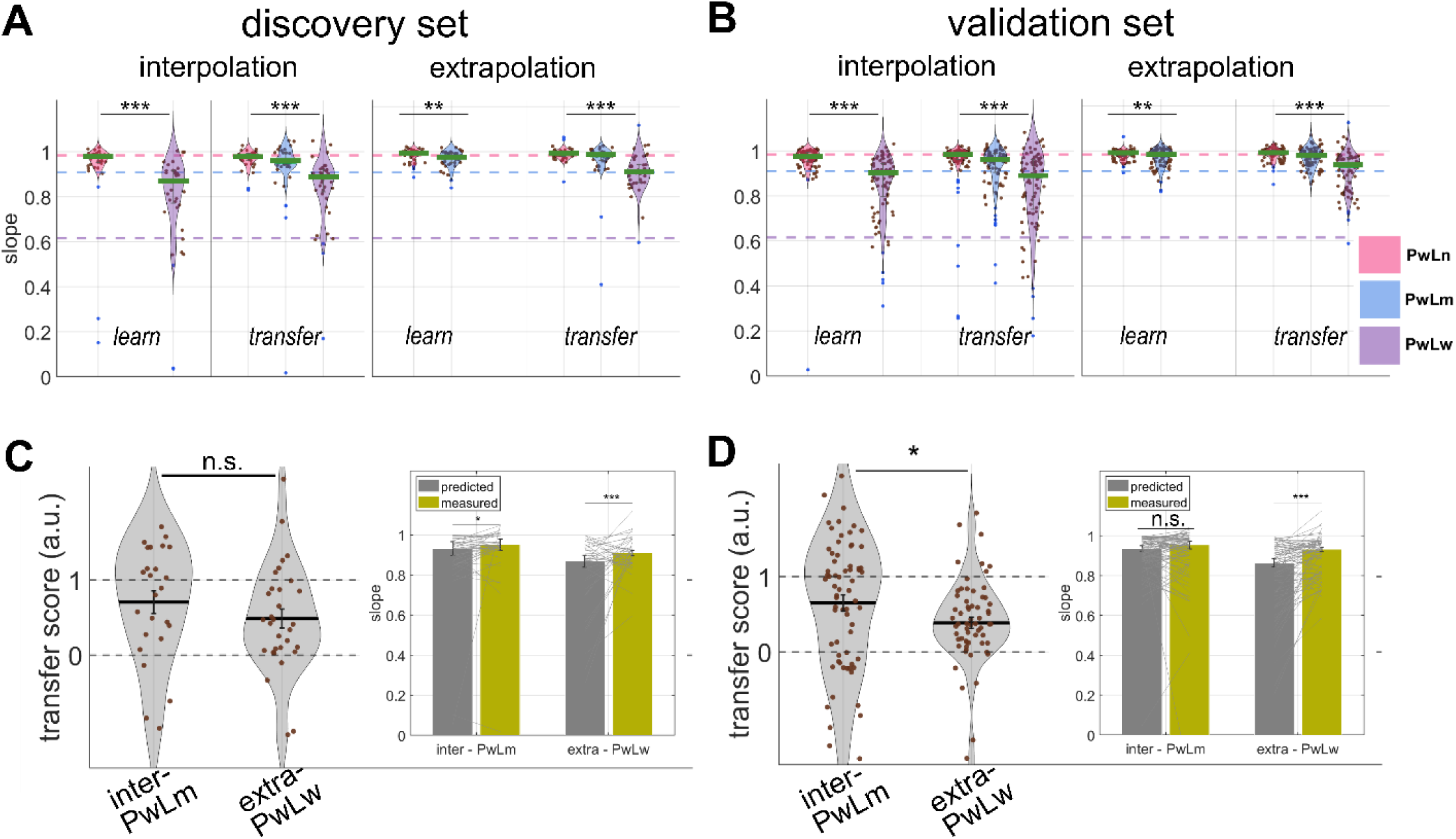
Slope and transfer score of experiment 2. A discovery & B validation set. Violin-plots show the distributions of slope in the learning and transfer phases of experiment 2, separated by likelihood and group. The central green line represents the median. The vertical bar within the violin spans the second and third quartiles. Each filled dot represents one participant. Three optimal slopes given likelihoods are presented as colour dashed lines, with pink representing PwLn, blue representing PwLm, and violet representing PwLw respectively. **C discovery & D validation set** Distributions of transfer scores in the interpolation and extrapolation groups. The central black line is the mean of each group, and the vertical bar is the standard error. Inlets showed predicted slope (gray bar) along with measured slope (olive bar) in the transfer-new trials. Note that there was no significant difference in the interpolation group, validation set. *p<= .05; ** p<= .01; ***p<=.001; n.s. non-significant

##### Validation set

A full model which includes the likelihood and group effects, and their interaction explained the slope data best (log likelihood ratio 13.82, *p* = .0001). A significant likelihood-group interaction was found (*p* = 1.74 X 10^-4^). Pos-Hoc analysis showed that slopes were qualitatively Bayesian for both interpolation (median ± iqr = .97 ± .06 for *PwLn* and median ± iqr = .89 ± .19 for *PwLw*; Wilcoxon sign rank test *p* = 3.73 X 10^-13^) and extrapolation (median ± iqr = .99 ± .02 for *PwLn* and median ± iqr = .98 ± .06 for *PwLm*; Wilcoxon sign rank test *p* = 3.25 X 10^-4^) groups, i.e. participants decreased sensory weighting as the variance of likelihood became bigger. We concluded that the interaction was caused by a larger disparity between prior/likelihood pairs in the interpolation group than which in the extrapolation group. There was an over reliance on likelihoods in the medium and wide likelihood conditions (right tailed sign rank test compared to optimal slopes; interpolation – *PwLw p* = 3.16 X 10^-13^, extrapolation – *PwLm p* = 1.33 X 10^-12^, **Fig 3.B**).

#### Transfer phase

##### Discovery set

We did not find significant slope differences between transfer-old trials and learning trials (median slope of “learning” trials = .97 & “transfer-old” trials = .98, Wilcoxon sign rank test *p* =.08). For all transfer trials, a likelihood-only model explained slope data best (log likelihood ratio as compared to the model with likelihood and group effects 3.96, *p* = .06). Slopes in the transfer phase remained Bayesian (likelihood effect *p* = 1.82 X 10^-08^).

Transfer scores (**Fig 4.C**) were significantly bigger than zero (right tailed t-test *p* = 1.15 X 10^-08^), implying the presence of transfer. The transfer score of the interpolation group was 0.70 ± .17 (Mean ± SE) while that of the extrapolation group was 0.58±.14 (Mean ± SE). Although the two group did not differ statistically (two sampled t-test *p* = 0.25), we identified one difference between the two groups: For the extrapolation group, the transfer score was significantly smaller than one (one sample t-test, *p* = 3.45 X 10^-04^). For the interpolation group, however, the transfer score only marginally differed from one (one sample t-test *p* = 0.06). Although the Bayesian factor (BF_01_) was 1.03, showing equivalent evidence supporting the null hypothesis.

##### Validation set

Comparing “old” trials between the learning and transfer phases, a model which includes the likelihood, group effects, and their interaction explained the slope best (log likelihood ratio as compared to a model including likelihood and group but no interaction 21.13, *p* < .001). It is noticeable that a model which also includes the phase effect failed to explain slope better than the winning model (log-likelihood ratio 2.98, *p* = .55), supporting no changes of slopes after removing position feedback. Indeed, the difference of slope values between transfer-old trials and learning trials was negligible (“learning” trials median ± iqr = .97 ± .07 versus “transfer-old” trials median ± iqr = .98 ± .06; Wilcoxon sign rank test *p* = .57). The post-hoc for a significant likelihood-group interaction (*p* = 4.09 X 10^-6^) found that the interaction was caused by a larger disparity between prior/likelihood pairs of the interpolation group (median ± iqr = .98 ± .04 for *PwLn*; median ± iqr = .88 ± .22 for *PwLw*; Wilcoxon sign rank test *p* = 1.25 X 10^-21^) than which of the extrapolation group (median ± iqr = .99 ± .03 for *PwLn*; median ± iqr = .98 ± .06 for *PwLm*; Wilcoxon sign rank test *p* = 1.36 X 10^-6^).

For all transfer trials, a model including the likelihood and group effects and their interaction explained the slope data better than any other models (log likelihood ratio 5.5, *p* = .01). There was a main likelihood effect (*p* = 3.16 X 10^-10^), indicating slopes in the transfer phase remained Bayesian-fashioned. Interaction was again found resulting from a larger disparity between each prior/likelihood pairs in the interpolation group than which in the extrapolation group (interpolation *PwLn – PwLm – PwLw* (median ± iqr) .98±.04 *–* .95±.13 *–* .87±.13 versus extrapolation *PwLn – PwLm – PwLw* (median ± iqr) .99±.03 *–* .98±.05 *–* .93±.15).

Transfer scores (**Fig 4.D**) were significantly higher than zero (right tailed t test *p* = 1.15 X 10^-08^), implying the presence of transfer. Transfer score (**Fig 4.D**) of the interpolation group was statistically larger than which of the extrapolation group (interpolation 0.65±.11 (Mean±SE); extrapolation 0.39±.17 (Mean±SE); two tailed t-test, *p* = .050).

### Optimality index

We compared the optimality index between the first and last 10 trials of transfer-new trials (**Fig 5**). There was no difference (paired t-test; **5.A** experiment 1-discovery *p* = .81, **5.B** experiment 1-validation *p* = .76, **5.C** experiment 2-discovery *p* = 0.92, **5.D** experiment 2-validation *p* = 0.09), showing that participants did not use partial feedback in the transfer phase to improve performance incrementally. Similarly, no difference was found between the first 10 transfer-old-trials and the last 10 in the learning phase (**Fig 5** right panel; paired t-test; **5.A** experiment 1-discovery *p* = 0.54, **5.B** experiment 1-validation *p* = 0.60 **5.C** experiment 2-discovery *p* = 0.83, **5.D** experiment 2-validation *p* = 0.63), further supporting that performance did not diminish after the removal of position feedback.

**Fig 5.**
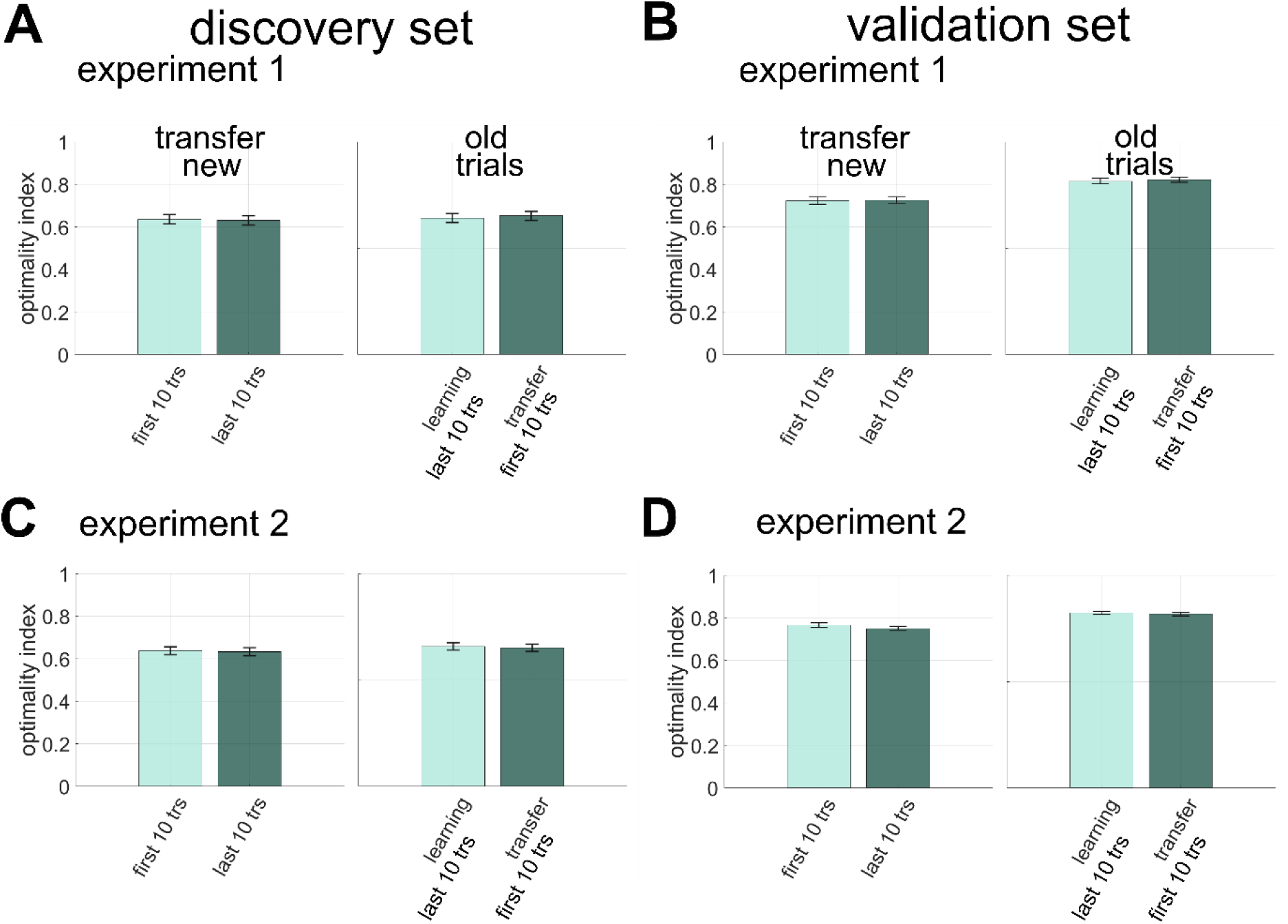
Optimality index. A discovery experiment 1 B validation experiment 1 C validation experiment 2 D validation experiment 2. Bar graphs show the mean optimality index and error bars represent standard errors. In the left panel of each subplot, the mean of first 10 trials of the transfer-new (light green bar) is compared against the mean of the last 10 trials (dark green bar). In the right panel, the mean of the last 10 trials of the learning phase (light green bar) is compared against the first 10 transfer-old trials (dark green bar).

## Discussion

The Bayesian decision model has achieved great success in describing human behaviours across various domains. However, whether humans make explicit Bayesian computation, i.e. learning and maintaining prior and likelihood distributions to compute posterior for decision, remains an ongoing debate. Examining whether people transfer experienced priors and likelihoods to a new scenario can help answer this question.

However, previous studies testing Bayesian transfer were inconclusive because they simply compared how close behaviours were to Bayes optimal in new scenarios. This is problematic because it largely depends on prior/likelihood estimation accuracy rather than transfer per se. Indeed, there are several reasons that can explain suboptimal behaviours, as previously discussed (32). Moreover, a direct quantification of transfer was lacking. To address these limitations, we devised transfer score – a mathematical marker that quantifies transfer without being hindered by prior/likelihood estimation biases. By applying the transfer score to the coin task (30), we found significant albeit suboptimal transfer in two perceptual decision studies. For each study, our results were replicated across two independent datasets (discovery and validation sets), suggesting robust replicability.

In the first study, we manipulated the way in which two location priors were learned and observed their effects on Bayesian transfer. The purpose was to better understand how much cognitive load can restrict the use of Bayesian computation. One argument against explicit Bayesian computation in human cognition is its complex calculations, and hence the greater demand it imposes on the brain compared to non-probabilistic computations (11,33). A previous study (22) tested Bayesian transfer when people concurrently learned two position priors. They found that weighting change in accordance with the newly introduced likelihood uncertainty did happen, but not sufficiently, and it only happened until an explicit instruction informing two different prior uncertainties was given. Thus, it was considered “suboptimal transfer”. We hypothesised if simultaneous multiple prior learning was the cause of such weak transfer they observed, sequentially presenting priors may rescue transfer performance as suggested by another study (20). We compared sequential (similar to (20)) and concurrent (similar to (22)) prior learning side by side. Our transfer scores showed that the two conditions led to equivalently suboptimal transfer. Thus, we did not find evidence supporting that simultaneous learning of two spatial priors prompted the use of non-Bayesian strategies to impede transfer. Interestingly, even though we explicitly instructed participants that our two priors had different levels of uncertainty, we could not identify transfer in the slope measurement as defined and observed in Kiryakova and colleagues (22). We concluded that these suboptimal behaviours found in past studies and ours are more likely a manifestation of imprecise uncertainty estimation rather than the execution of non-probabilistic computation.

The critical question is how much people can generalize knowledge to unseen data in any given task. We addressed the question head on in the second study. We compared the transfer score between generalization within (interpolation) and outside (extrapolation) the range of experienced sensory information (likelihoods) when performing our visuospatial task. Again, there was significant but suboptimal transfer in both conditions. Between the two conditions, people had a lower transfer score in the extrapolation and this difference was statistically significant in the (larger) validation dataset. These results indicated that generalization is better when new data points are within the context of past observations. This is consistent with what Kiryakova and colleagues (22) have shown. However, our design shielded any observed effect from the influence of imperfect uncertainty estimation (in the approximation algorithm) and is hence more robust. Moreover, unlike previous studies, we quantified the degree of transfer.

Taking findings of the two experiments together, we ask, what is the most possible computation strategy for the coin task? There was no evidence supporting the use of a heuristic strategy in the experiment 1. A stable optimality index also suggests participants did not make decisions using model-free (error-based) learning. A Bayesian observer that acquires a generative model of the task, however, would have behaved similarly between interpolation and extrapolation, which was not what we observed in the second experiment. The results of the two experiments seemed to be at odds, with the first supporting and the second casting a doubt on Bayesian computation, albeit with knowledge transfer. It is possible that participants acquired a probabilistic recognition model linking contextualised sensory inputs to corresponding policies (17,34). The model is optimised by training data to approximate the true generative model. It would generalize well enough within, but not outside, past experience because it does not represent the true generative process (**Fig 6.A**). However, we cannot fully rule out that participants capitalised on the feature of the coin task to behave in a Bayesian fashion using a special case of heuristics (35). Recall that the optimal policy is a linear mapping between stimuli and response (eq 1). Participants could have performed well enough by learning one mapping for one stimulus type without acquiring any prior or likelihood information. They could then combine previously learned mapping in the transfer trials to mimic generalization. One might think that a linear-mapping learner should paradoxically generalize equally well in interpolation and extrapolation. We argue that due to the biased estimates observed in our data, it is only reasonable to see lower transfer in extrapolation if our participants used a linear mapping policy (**Fig 6.B**).

**Fig 6.**
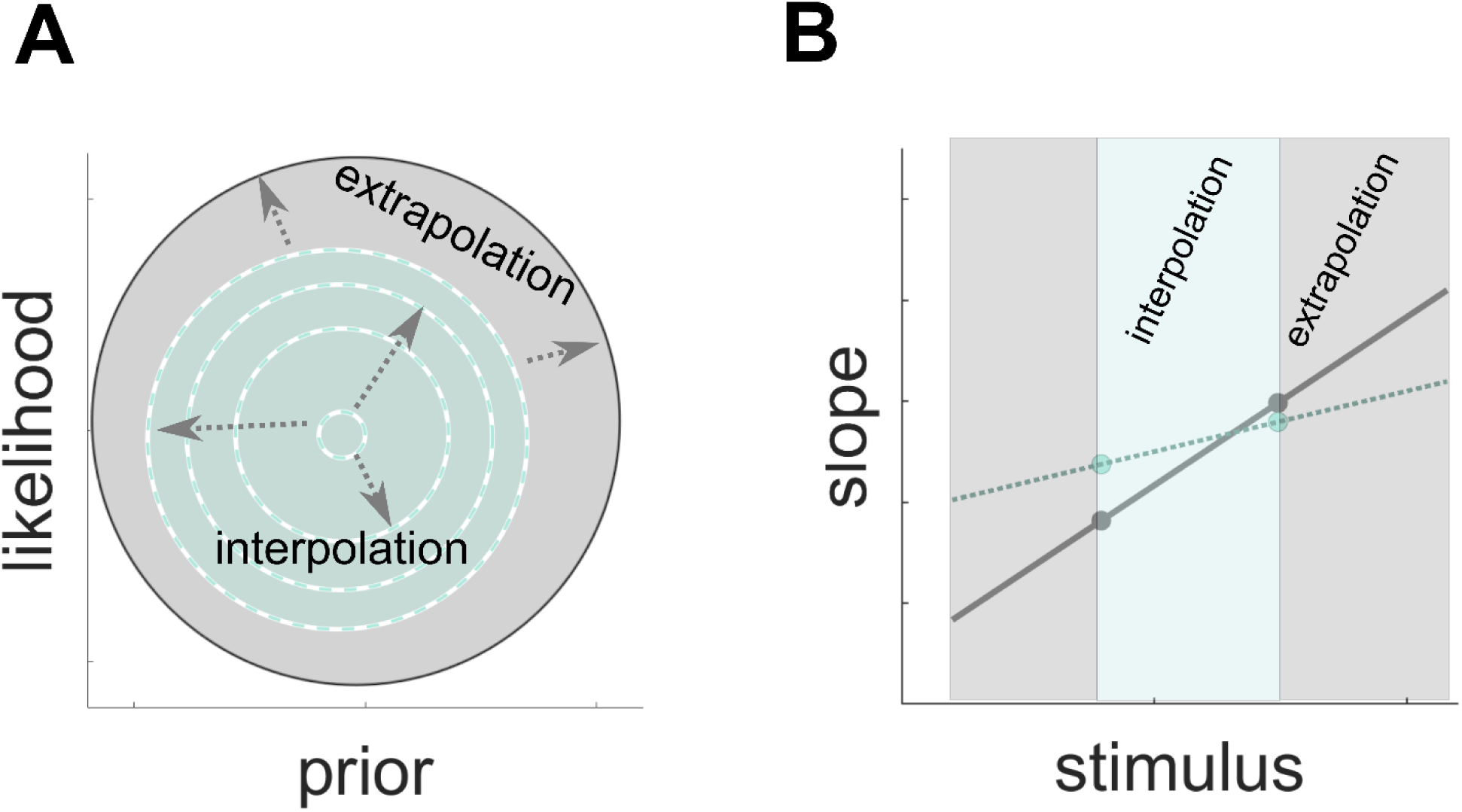
**A** schematic illustration of a probabilistic recognition model which approximates the true generative model (grey circle). The true posterior of is a function of prior (x-axis) and likelihood(y-axis). Dashed circles represent experienced data. The approximate posterior matches the true posterior better in the area that is within (green area) rather than outside experience (grey area). Figure adapted from (17,18) **B** schematic illustration of a possible linear mapping strategy for the coin task and how it could result in behaviours similar to our findings in the experiment 2. Grey dots represent Bayes optimal slopes (y-axis) given stimuli (x-axis). An optimal linear mapping between scatters of splashes and slopes (grey line) will produce optimal extrapolation even if no true generative process is represented. In reality, because of noisy slope estimations (green dots), interpolation (green zone) would be closer to optimal than extrapolation (grey zone).

Our studies nevertheless showed that people may not perform decisions under uncertainty by computing the full generative model. This is in line with many neurocomputational theories formulating that approximation algorithms are needed for human Bayesian cognition (17,36–38). We showed that perceptual decisions use Bayesian approximation with knowledge transfer, albeit suboptimal. We also demonstrated that we can more effectively validate transfer using a rigorous mathematical definition of transfer score, that can be applied to various probabilistic inference tasks and advance our computational understanding of human behaviours. It would be especially valuable to test transfer in a similar perceptual decision task that uses likelihoods presenting realistic noise (such as luminance). Based on the abovementioned bounded-optimal hypothesis, transfer will be close to optimal in both interpolation and extrapolation case given that our brain has been trained by such types of likelihood through life-long experience (i.e., hardwired priors) while linear mapping learners are likely to behave similarly to the current study.

More studies are needed to confirm the exact computation underpinning decisions. Recently, another paradigm (39) was proposed for scrutinising whether humans make inference using explicit Bayesian computation. The paradigm uses multiple prior-likelihood pairs in the Bayesian decision model which lead to an identical decision policy and investigates the learning dynamics when new component (i.e. prior-likelihood) pairs being introduced to discriminate explicit Bayesian learner from policy learner. In short, right after a transition to a new pair, explicit Bayesian learners’ performance would temporarily deteriorate as they re-learn the new prior/likelihood while policy learners would maintain the same level of performance. We previously argued that some tasks are more sensitive to transfer manipulations while others are more sensitive to component-pair manipulations and the two can be complementary in validating true Bayesian learners behaviourally (32). Future work could make use of these two complementing paradigms to systematically inquiring the exact process model that supports different types of decision under uncertainty.

## Methods

### Participants

The online research was approved by the University of Melbourne research ethics committee (research ethics 2021-20592-18973-4). Participants were recruited using the University of Melbourne psychology research participation pool and Prolific online survey platform (prolific.co). All participants completed self-reported questionnaire to confirm that they had normal or corrected-to-normal visual acuity, and no history of neurological, psychiatric disorders or substance use, and gave written informed consent prior to taking part in the study. Participants were compensated with course credits or £5 per hour for their time.

For experiment 1, 102 adults (74 females, age mean ± SD = 19.81 ± 3.92 yrs) were recruited for the discovery study. Among these participants, data from 6 participants were excluded entirely, 1 for not meeting the inclusion criteria and 5 for poor data quality (exclusion criterion based on data quality see below **Data analysis**), resulting in a final sample of 96 participants (48 for each group). For the validation study, 170 participants (120 females, age mean ± SD = 19.90 ± 5.60 yrs) were recruited. Among them, 11 participants were excluded for poor data quality, resulting in a final sample of 159 participants (80 for serial group and 79 for parallel group). There was neither significant age nor gender difference between the discovery and validation set.

For experiment 2, 99 participants (66 females, age mean ± SD = 20.36 ± 3.91 yrs) were recruited for the discovery study. Among these participants, data from 13 participants were excluded for history of neurological or psychiatric disorders and 11 for poor data quality, resulting in a final sample of 75 participants (35 for interpolation and 40 for extrapolation group). For the validation study, 174 participants (136 females, age mean ± SD = 19.71 ± 3.28 yrs) were recruited. Among them 10 participants were excluded for poor data quality, resulting in a final sample of 164 participants (82 for each group). There was neither significant age nor gender difference between the discovery and validation set.

Our power analysis was based on the findings of the discovery data (R software: “SSDbain” and Fu et al., 2021), indicating that, to increase the power to 80% to confirm a moderate effects at *a* = 0.05 for a one-sided t test, 80 participants would be required for each group.

## Experiments

All experiments were designed using PsychoPy (PsychoPy 2020) and launched online through the Pavlovia platform. Stimuli were displayed on participants’ own screens. The minimum requirement for a screen resolution was 1172×553 pixel.

### Coin Task

All stimuli were in an arbitrary unit that defines the horizontal location of the left and right edges of the screen as -0.5 and 0.5. Participants were instructed to view the screen as the surface of a pond and to locate an unseen coin that a person had thrown into the water (**Fig 1.A**). At the beginning of each trial, participants saw five dots (diameter = 0.01) which represent the “splashes” caused by a coin dropping to the pond. The horizontal positions of these five dots were drawn from a Gaussian “likelihood” distribution with a mean of the horizontal coin position in that trial and a standard deviation assigned from one of the three values, (***σ_L_*** = 0.0024, 0.06 and 0.15). In each trial, the horizontal location of the coin was drawn from a Gaussian “prior” distribution which centres at the middle of the screen (mean = 0) and has a standard deviation ***σ_p_*** of either 0.025 or 0.085 (**Fig 1.B**). Participants’ task was to catch the coin by moving a vertical blue bar (the “net”, width ***l*** = 0.02) to their estimated hidden coin position and then click on the “space” button to indicate the answer. As the height of the bar equalled the height of the screen, vertical locations of splashes and coin made no difference to participants. There was no temporal deadline, so participants had all the time they needed to submit their response. After responding, participants received feedback information for one second before the next trial started automatically. There were two phases: learning and transfer phases, in the coin tasks (see more information about the two phases in **Methods - Experiment specific details**). Feedback information of the two phases differed as follows (**Fig 1.A** **lower panel)**. In the learning phase, participants received trial-by-trial coin location feedback, which was displayed as one yellow dot (diameter ***d*** = 0.01) along with splashes. If there was an overlap between the net and the coin, this trial was deemed a correct trial and participants’ scores increased by one point. Coin location feedback was given in every trial and for every successful trial the accumulative score was displayed along with it. In the transfer phase, only scores but not coin positions were given. There were minor differences in how the scores were shown between the discovery and validation study during the transfer phase. In the discovery set, scores were shown in all corrective trials. In the validation set, participants received a summary of their total correct trials every 15 trials during the transfer phase. In both sets, at the beginning of the transfer phase, we informed participants about this change of feedback. However, we also reminded participants that throughout the experiment the goal was to be as accurate as possible in locating the coin, irrespective of feedback.

### Likelihood-only task

We also administered a likelihood-only task to estimate perceived likelihood uncertainty. In each trial, participants saw five “splashes” on the screen. They were instructed to move the “net” to where they thought the “centre” of the 5 dots on the horizontal axis (x-axis) was. After responding, the true horizontal location of the “centre” was revealed by an orange vertical bar (**Fig 1.A** left **lower panel**) which had a height equalling the height of the screen and a width .006. Previously (41) it was found that estimations of splash centres were biased by priors when people undertook the likelihood-only task after the coin task. Therefore, we administered the likelihood-only task before the coin task in the experiments.

### Visual memory task

In both experiments, a Corsi block-tapping test (42) was used to assess participants’ working memory before they started the likelihood-only task. The task began by flashing several blocks on the screen. Participants were required to respond by tapping the block either in the forward or backward order of flashing. The test started with three flashes, the number of flashes increased by one after a participant gave one correct answer and decreased by one after a participant gave two successive incorrect answers. There were 16 working-memory trials in total, eight in a block of serial order followed by eight trials in a block of reverse-serial order. We used this task as a screening task. Participants who failed to correctly respond to 3 flashes were excluded from further analysis.

### Experiment 1 specific details (Fig 1.C)

Participants were randomly assigned to a serial or parallel learning group. The two groups only differed in the presentation order of trial types (prior-likelihood pairs) in the learning phase. Both groups completed the likelihood-only task, followed by the learning, and then transfer phases of the coin task. The likelihood-only task constituted of total 80 trials, interspersing 40 of each likelihood distribution (narrow ***σ_Ln_*** = 0.06 and wide ***σ_Lw_*** = 0.15). The learning phase of the coin task constituted of 400 trials, 200 trials of two sets of prior (narrow ***σ_Pn_*** = 0.025 and wide ***σ_Pw_*** = 0.085) and likelihood (narrow ***σ_Ln_*** = 0.06 or wide ***σ_Lw_*** = 0.15) combinations. Overall, there were four possible prior-likelihood combinations, i.e. *PnLn, PnLw, PwLn* and *PwLw*. For each participant, the combinations were always orthogonalized such that one likelihood only paired with one prior but not the other during learning. Prior-likelihood combinations were counterbalanced across participants. For the serial learning group, participants learned one combination in trial 1-200 (4 blocks of 50 trials) and then the other in trial 201-400. For the parallel learning group, the two prior-likelihood pairs were interleaved throughout the leaning phase. We used a graph instruction to inform participants that there are two throwers with different levels of accuracy hitting a coin at every block beginning. During a trial, throwers were represented by two splash colours – green and blue, with the colour and prior pairing counterbalanced across participants. In the transfer phase, both groups experienced 180 trials interspersing 90 of each likelihood distribution paring with one prior.

### Experiment 2 specific details

Participants completed the likelihood-only task, followed by the learning, and then transfer phases of the coin task. In the likelihood-only task, there were 90 trials with 30 trials of each likelihood distribution (narrow ***σ_Ln_*** = 0.024, medium ***σ_Lm_*** = 0.06, and wide ***σ_Lw_*** = 0.15) interspersing over three 30-trial blocks. Only one prior (***σ_Pw_*** = 0.085) was used for the experiment 2.

Participants were randomly assigned to an “interpolation” or “extrapolation” group for the coin task (**Fig 1.D**). For the interpolation group, participants were presented with 100 *PwLn* and 100 *PwLw* trials in the learning phase. For the extrapolation group, participants experienced 100 *PwLn* and 100 *PwLm* trials in the learning phase (**Fig 1 D**). During the transfer phase, all participants encountered *PwLn*, *PwLm* and *PwLw* trials pseudorandomly (225 trials, 75 trials for each prior/likelihood pair). Therefore, for the interpolation group the new combination in the transfer task was *PwLm* while for the extrapolation group was *PwLw* trials.

### Data analysis

All the data analysis were performed using R studio (R 4.1.1) and Matlab (Mathworks release 2021b). Data exclusion was based a median absolute deviation (MAD) to identify and exclude outliers in trials and individual participants (43). It would exclude any data points that was 3 MAD away from the median. For grouped analysis, the MAD criterion was first used to identified outliners in each prior/likelihood combination. Participants presenting with outlier slopes (see below) in all combinations used for the transfer phase were excluded from grouped analyses.

### Slope: estimating likelihood reliance

According to the Bayes rule, an optimal estimation of the coin location (*X_est_*) is:

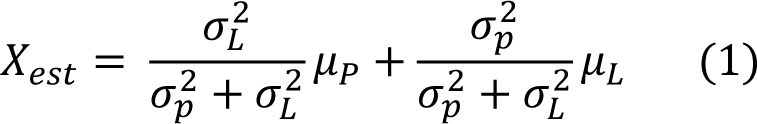

That is, an optimal estimated position is a weighted average of prior mean *μ_P_* and likelihood mean *μ_L_* and the weighting is based on the relative precision (= the inverse of variance *σ^2^*), with 𝜎𝜎^2^ being the prior variance and *σ^2^* being the variance associated with the likelihood, which can be estimated by *σ^2^* = likelihood variance/number of dots (5 in our case). We can compare a participant’s likelihood weight against this optimal decision to learn if behaviours are optimal. Similar to (44), for each prior/likelihood combination in each participant, we used the polyfit.m function in Matlab to fit chosen net locations *x_net_* a linear function of cue centre positions *Xnet* = *intercept* + β μ*L*. The slope of the regression line (= 𝛽) indicates how much participants relied on likelihood, i.e. sensory weight (**Fig 2.A**), and is expected to equal 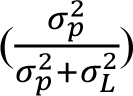 for a Bayesian optimal observer. The closer the slope is to one, the more a participant relies on the 𝜎 +𝜎 𝑝 𝐿 likelihood. The closer the slope is to zero, the more a participant relies on the prior.

### Predicted slope in the transfer phase based on subjective prior and likelihood

While people changed slopes according to the reliabilities of priors and likelihoods in previous coin task studies, it has been shown that they often placed more weights on the likelihood than ideal Bayes observers would have (22,45). One very possible explanation is that while researchers expect observers place weighting according to objective uncertainty, i.e. experimenter-designed values, people tend to weight based on subjective uncertainty (22,41). Therefore, a slope value which deviates from Bayes optimal during transfer may be a manifestation of imperfect uncertainty estimation, a result of a non-Bayesian strategy or both. Simply measuring slopes cannot discriminate the two. Our solution is to compare measured transfer-phase slope against predicted transfer-phase slope based on subjective prior and likelihood estimations. To do so, first, we estimated participants’ subjective likelihood variance (𝜎^2^_*Lsub*_) as follows

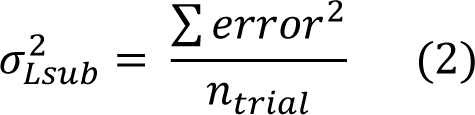

Here an error on each likelihood-only trial was calculated by taking the difference between the response and true centre for that trial. Second, we computed the subjective prior variance (41,46) such that

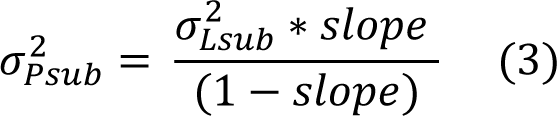

Third, subjective *σ^2^_Psub_* and *σ^2^_Lsub_* were plugged into (eq 1) to acquire a predicted transfer-phase slope for each participant shall they perfectly combine subjective prior and likelihood uncertainties during transfer.

### Transfer score (Fig 2.B)

Expanding on the predicted transfer-phase slope, we developed a normalised measure, called the transfer score, to quantify how well people generalize knowledge about prior uncertainty that they acquired in the learning phase into the transfer phase. As abovementioned, a predicted slope using subjective estimates of prior and likelihood is computable so it can be compared against a measured change. A normalised transfer score (*ts*) thus is defined as follows:

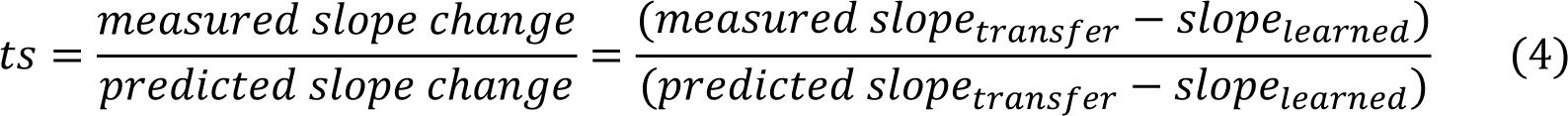

Note that the transfer score is a directional measure with a score of one meaning optimal transfer, a score equal to or smaller than zero meaning no transfer, and a score between zero to one indicating suboptimal transfer. In the experiment 2, for each participant we would have acquired two measured slope changes using the two prior/likelihood combinations during the learning phase. We took their mean to calculate the transfer score.

### Optimality index (Fig 2.C)

To understand how behaviours changed dynamically during the experiment, we adapted “optimality index” from (21) to quantify participants’ performance on a trial-by-trial basis. The rationale of optimally index is as follows. For every horizontal position on the screen ***x***, we can calculate the probability of hitting the true coin location *p_hit_* given a participant placing the net on ***x***. The analytical solution of *p_hit_* based on the mean ***μ*** *and standard deviation of **σ*** of the true Gaussian posterior for each prior and likelihood combination is

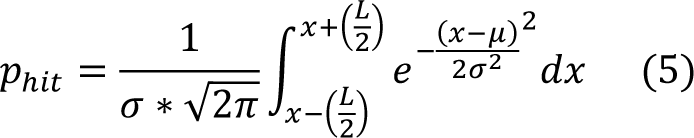

***L*** equals the width of the net ***l*** plus the diameter of the coin ***d.*** *p_hit_* is visualised in the **Fig 2.C** as the area under the probability density function curve. The hitting probability is maximal (= *max(p_hit_)*, illustrated as the yellow area in the **Fig 2.C**) when ***x*** equals ***μ***. The optimality index for a given trial is thus defined as *p_hit_(x_net_)/max(p_hit_)*.

### Statistical analysis

We used the Kolmogorov-Smirnov test to evaluate the normality of data. Parametric tests including mixed-design ANOVA and t-test were used for normally distributed data. For data which violated the assumption of normality we used fitlme.m in Matlab to perform a linear mixed effects analysis and Wilcoxon tests to compare the medians. The significance level of all tests was 0.05. Where appropriate, Bayes factors were also reported in support of evidence for the null hypothesis.

## Acknowledgements

We are grateful to Yuanyuan Li and Sara Chen for their help with recruiting and running participants.

## Author Contributions

**Conceptualization:** Chin-Hsuan Sophie Lin, Marta I Garrido

**Formal analysis:** Chin-Hsuan Sophie Lin, Trang Thuy Do, Lee Unsworth

**Funding acquisition:** Chin-Hsuan Sophie Lin, Marta I Garrido

**Investigation:** Trang Thuy Do, Lee Unsworth

**Methodology:** Chin-Hsuan Sophie Lin, Trang Thuy Do, Lee Unsworth

**Project administration:** Chin-Hsuan Sophie Lin, Marta I Garrido

**Supervision:** Chin-Hsuan Sophie Lin, Marta I Garrido

**Validation:** Chin-Hsuan Sophie Lin

**Writing – original draft:** Chin-Hsuan Sophie Lin, Marta I Garrido

**Writing – review & editing:** Chin-Hsuan Sophie Lin, Trang Thuy Do, Lee Unsworth, Marta I Garrido

## Supporting information

**S1 Fig. Experiment 1 transfer score for each prior/likelihood combination, separated by cognitive load group**

