## supplemental figure 1 for "Are we really Bayesian? Probabilistic inference shows sub-optimal knowledge transfer"

### Supplementary figures

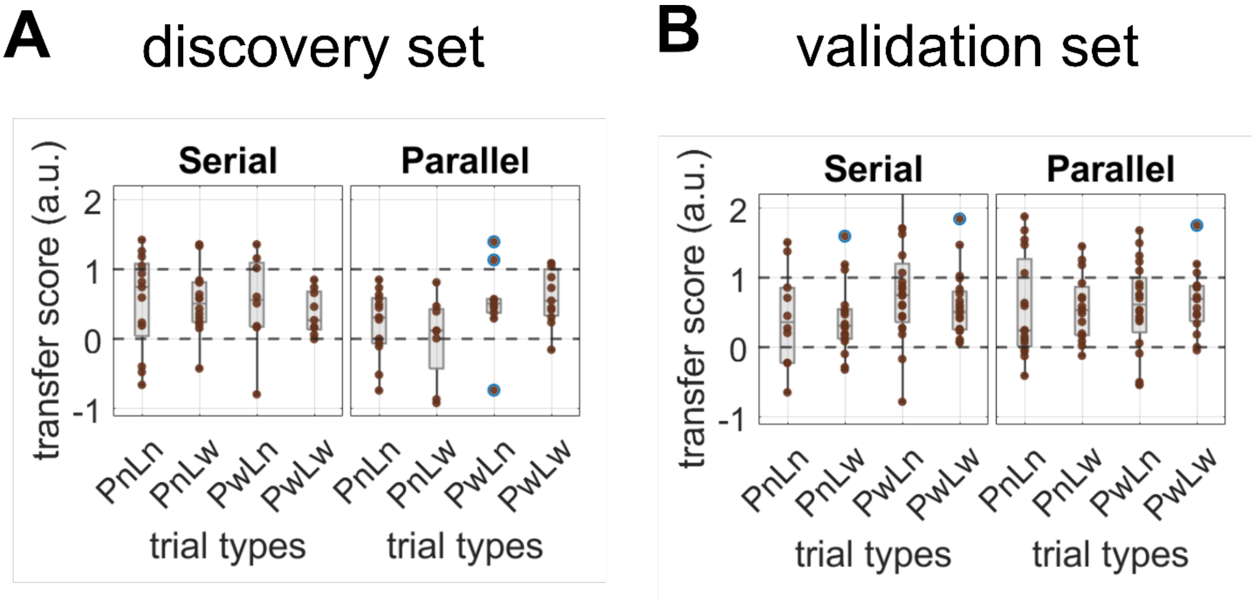


***S1 Fig.*** *Experiment 1 transfer score for each prior/likelihood combination, separated by cognitive load group* ***A discovery set B validation set.***
